# Dynamic load balancing enables large-scale flux variability analysis

**DOI:** 10.1101/440701

**Authors:** Marouen Ben Guebila

## Abstract

Genome-scale metabolic models (GSMMs) of living organisms are used in a wide variety of applications pertaining to health and bioengineering. They are formulated as linear programs (LP) that are often under-determined. Flux Variability Analysis (FVA) characterizes the alternate optimal solution (AOS) space enabling thereby the assessment of the robustness of the solution. fastFVA (FFVA), the C implementation of MATLAB FVA, allowed to gain substantial speed up, although, the parallelism was managed through MATLAB. Here veryfastFVA (VFFVA) is presented, which is a pure C implementation of FVA, that relies on lower level management of parallelism through a hybrid MPI/OpenMP. The flexibility of VFFVA allowed to gain a threefold speedup factor and to decrease memory usage 14 fold in comparison to FFVA. Finally, VFFVA allows processing a higher number of GSMMs in faster times accelerating thereby biomedical modeling and simulation.
VFFVA is available online at https://github.com/marouenbg/VFFVA.

---

Modeling and simulation of biological systems gained tremendous interest thanks to the increasing predictive ability of the modeled systems in healthcare and the biotechnology industry [6]. Microbial and human systems are most amenable to modeling given the development of high-throughput techniques that enable the spatiotemporal characterization of biological systems [18].

Particularly, constraint-based reconstruction and analysis (COBRA) methods enable the reconstruction of the metabolism of biological systems *in silico* as linear programs [19]. Subsequently, an objective function of the system is formulated and optimized for, e.g., biomass yield, metabolite production. Although the objective is uniquely determined, the set of corresponding solutions forms the space of alternate optimal solutions (AOS) that describe the possible conditions in which the optimal objective is achievable. The AOS space is quantified using flux variability analysis (FVA) [11], which provides a range of minimum and maximum values for each variable of the system. Biologically, these values overlap with the fitness of a given system to achieve optimality and allow to validate the metabolic phenotype through matching the empirical ranges with the FVA bounds. FVA was applied to quantify the fitness of macrophages after the infection of *Mycobacteirum tuberculosis* [2], resolve thermodynamically infeasible loops [14], and compute the essentiality of reactions [4]. fastFVA (FFVA) [7], a recent implementation of FVA gained tremendously in speed over the `fluxvariability` COBRA toolbox MATLAB function [10]. Two main improvements were the driving factor of the gained efficiency: first, the C implementation of FVA allowed higher flexibility in comparison to MATLAB [12] through the use of the CPLEX C API. The second was the use of the same LP object, which avoided solving the problem from scratch in every iteration, thereby saving presolve time. FFVA is compiled as MATLAB Executable (MEX) file, that can be called from MATLAB directly.

Nevertheless, given the exponentially growing size of metabolic models, FFVA is run in parallel in most cases. Parallelism simply relies on allocating the cores through MATLAB parpool function [12] and running the iterations through parfor loop. The load is statically balanced over the workers such as they process an equal amount of iterations. Nevertheless, the solution time varies greatly between LPs which does not guarantee an equal processing time among the workers in static load balancing. Often, the workers that were assigned a set of fast-solving LPs process their chunk of iterations and stay idle, waiting to synchronize with the remaining slower workers, which can result in less efficient global run times. Here I present veryfastFVA (VFFVA), which is a pure C implementation of FVA, that has a lower level management of parallelism over FFVA. The program is provided as a standalone, and does not rely on MATLAB thereby offering an open source alternative for constraint-based biological analysis. The significant contribution lies in the management of parallelism through a hybrid OpenMP/MPI, for shared memory and non-shared memory systems respectively, which offers excellent flexibility and speed up over the existing implementations. While keeping the up-mentioned advantages of FFVA, load balancing in VFFVA was scheduled dynamically [20] in a way to guarantee equal run times between the workers. The input does not rely on MATLAB anymore as the LP problem is read in the industry standard *.mps* file, that can also be obtained from the classical *.mat* files through a provided converter. The improvements in the implementation allowed to speed up the analysis by a factor of three and reduced memory requirements 14 fold in comparison to FFVA and the Julia-based distributedFBA implementation [9], in a similar parallel setting.

Taken together, as metabolic models are steadily growing in number and complexity, their analysis requires the design of efficient tools. VFFVA allows making the most of modern machines specifications to run a more considerable amount of simulation in less time thereby enabling biological discovery.

## Material and methods

### Flux variability analysis

The LP problem modeling the metabolism of a given organism has *n* reactions that are bounded by lower bound *lb*_(*n*,1)_ and upper bound *ub*_(*n*,1)_ vectors. The matrix *S* _(*m*,*n*)_ represents the stoichiometric coefficients of each of the *m* metabolites involved in the *n* reactions. The system is usually constrained by *S*.*v* = 0 to represent the steady-state, also referred to as Flux Balance Analysis (FBA) [17]. An initial LP optimizes for the objective function of the system to obtain a unique optimum, e.g., biomass maximization, like the following:

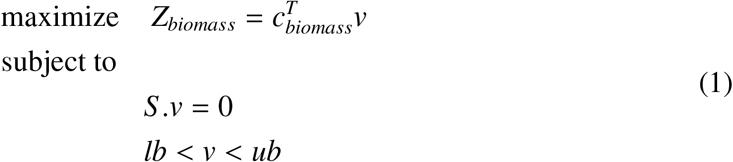

The system being under-determined (*m* < *n*), there can be an infinity of solution vectors *V*_(*n*,1)_ that satisfy the unique optimal objective (*c*^*T*^*v*), with *C*_(*n*,1)_ as the objective coefficient vector. In a second step, in order to delineate the AOS space, the objective function is set to its optimal value followed by an iteration over the n dimensions of the problem. Consequently, each of the reactions is set as a new objective function to maximize (minimize) the LP and obtain the maximal (minimal) value of the reaction range. The total number of LPs is then equal to 2*n* in the second step which is described as the following:

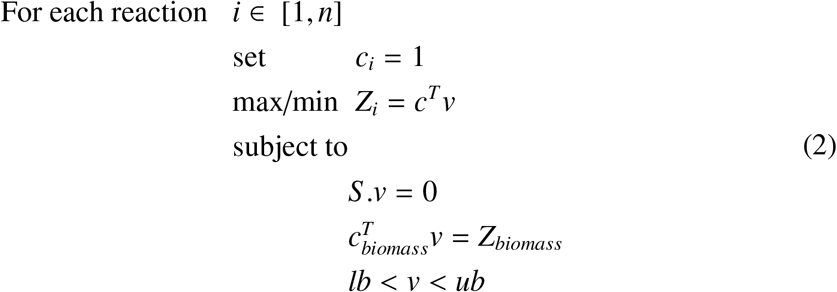

The obtained minimum and maximum objective value for each dimension define the range of optimal solutions.

### Management of parallelism

Problem 2 is entirely parallelizable through distributing the 2*n* LPs among the available workers. The strategy used so far in the existing implementations was to divide 2*n* equally among the workers. Nevertheless, the solution time can vary widely between LPs because ill-conditioned LPs can induce numerical instabilities requiring longer solution times. Consequently, dividing equally the LPs among the workers does not ensure an equal load on each worker.

Since it is challenging to estimate *a priori* the run time of an LP, the load has to be dynamically balanced during the execution of the program.

In shared memory systems, Open Multi-Processing (OpenMP) library allows balancing the load among the threads dynamically such that every instruction runs for an equal amount of time. The load is adjusted dynamically, depending on the chunks of the problem processed by every thread. At the beginning of the process, the scheduler will divide the original problem in chunks and will assign the workers a chunk of iterations to process. Each worker that completes the assigned chunk will receive a new one until all the LPs are processed.

In systems that do not share memory, Message Passing Interface (MPI) was used to create instances of Problem 2. Every process then calls the shared memory execution through OpenMP.

In the end, the final program is comprised of a hybrid MPI/OpenMP implementation of parallelism which allows great flexibility of usage, particularly in High-Performance Computing (HPC) setting.

### Another application: generation of warmup points

The uniform sampling of metabolic models is a common unbiased tool to characterize the solution space and determine the flux distribution per reaction [3, 13]. Sampling starts from pre-computed solutions called warmup points, from which the sampling chains start exploring the solution space. The generation of *p* ≥ 2*n* warmup points is done similarly to FVA. The first 2*n* points are solutions of the FVA problem, while the points ≥ 2*n* are solutions corresponding to a randomly generated coefficient vector *c*. Another difference with FVA lies in the storage of the solutions *v* rather than the optimal objective *c*^*T*^*v*. The generation of 30,000 warmup points was compared using the COBRA toolbox function createWarmup_MATLAB_ and a dynamically load-balanced C implementation createWarmup_VF_ that was based on VFFVA.

### Model description

A selection of models [7] was tested on FFVA and VFFVA. The models (Table 1) are characterized by the dimensions of the stoichiometric matrix *S*_*m*,*n*_. Each model represents the metabolism of human or bacterial systems. Models pertaining to the same biological system with different *S* matrix size, have different levels of granularity and biological complexity.

**Table 1:** Model size and description.

| Model | Organism | Size |
| --- | --- | --- |
| Ecoli_core [16] | *Escherischia coli* | (72,95) |
| P_putida [15] | *Pseudomonas putida* | (911,1060) |
| EcoliK12 [5] | *Escherischia coli* | (1668,2382) |
| Recon2 [23] | *Homo sapiens* | (4036,7324) |
| E_Matrix [21] | *Escherischia coli* | (11991,13694) |
| E<sub>c</sub>_Matrix [22] | *Escherischia coli* | (13047,13726) |
| Harvey [24] | *Homo sapiens* | (157056,80016) |

### Hardware and software

VFFVA and createWarmup_VF_ were run on a Dell HPC machine with 72 Intel Xeon E5 2.3GHz cores and 768 GigaBytes of memory. The current implementation was tested with Open MPI v1.10.3, OpenMP 3.1, GCC 4.7.3 and IBM ILOG CPLEX academic version (12.6.3). FFVA and createWarmup_MATLAB_ were tested with MATLAB 2014b [12] and dis-tributedFBA was run on Julia v0.5. ILOG CPLEX was called with the following parameters:

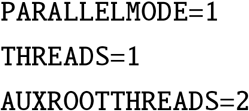

Additionally, large-scale coupled models with scaling infeasibilites might require

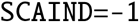

The call to VFFVA is done from bash as follows:

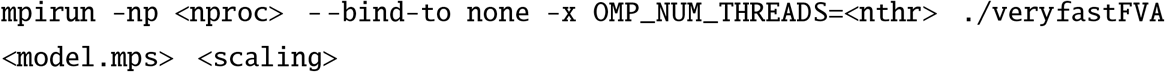
with *nproc* is the number of non-shared memory processes, *nthr* is the number of shared memory threads, *scaling* is CPLEX scaling parameter where 0 leaves it to the default (equilibration) and −1 sets it to unscaling. createWarmup_VF_ was called in a similar fashion: 
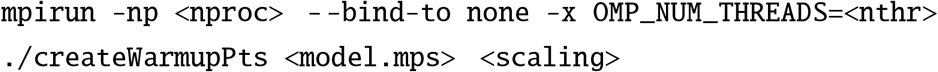

For large models, OpenMP threads were bound to physical cores through setting the environment variable

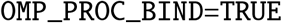

while for small models, setting the variable to `FALSE` yielded faster run times. The schedule is set through the environment variable

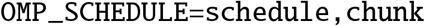

where `schedule` can be `static`, `dynamic` or `guided`, and `chunk` is the minimal number of iterations processed per worker at a time. The source code is available online [8].

### Other possible implementations

The presented software can be implemented in Fortran since the library OpenMP is supported as well. Additionally, Python’s multiprocessing library allows to balance the load dynamically between non-shared memory processes, but the parallelism inside one process is often limited to one thread by the Global Interpreter Lock (GIL). This limitation could be circumvented through using OpenMP and Cython [1]. The advantage of VFFVA lies in the implementation of two levels of parallelism following a hierarchical model where MPI processes are at a top-level and OpenMP threads at a lower level. The MPI processes manage the coarse-grained parallelism, and OpenMP threads manage the finer-grained tasks that share memory and avoid copying the original problem, which increases performance and saves consequent memory. This architecture adapts very well with modern distributed hardware in HPC setting.

## Results

The OpenMP/MPI hybrid implementation of VFFVA allowed gaining important speedup factors over the static load balancing in the MATLAB implementation. In this section, the run times of VFFVA were compared to FFVA at different settings then the different strategies of load balancing were compared through their impact on the run time per worker. While in FFVA the authors benchmarked serial runs [7], in the present work, the emphasis was put upon parallel run times.

### Parallel construct in a hybrid OpenMP/MPI setting

The MATLAB implementation of parallelism through the parallel computing toolbox provides great ease-of-use, wherein two commands only are required to allocate and launch parallel jobs. Also, it saves the user the burden of finding out if the jobs are run on memory sharing systems or not. VFFVA provides the user with a similar level of flexibility as it supports both types of systems and ensures sensibly the same numerical results as FVA. Besides, it allows accessing advanced features of OpenMP and MPI such as dynamic load balancing. The algorithm starts first by assigning chunks of iterations to every CPU (Figure 1), where a user-defined number of threads simultaneously processes the iterations. In the end, the CPUs synchronize and pass the result vector to the master core to reduce them to the final vector.

**Figure 1:**
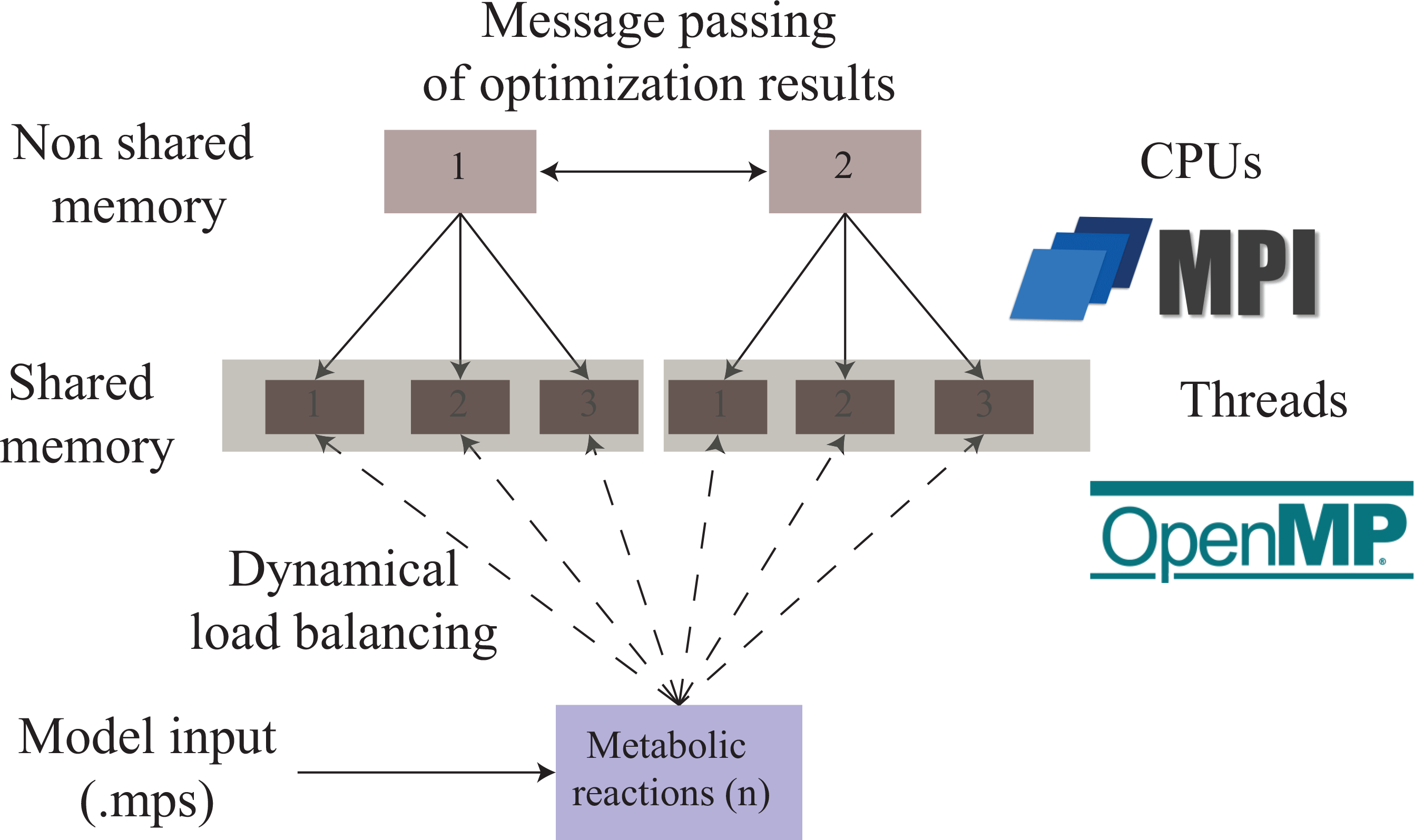
Hybrid OpenMP/MPI implementation of FVA ensures two levels of parallelism. The distribution of tasks is implemented following a hierarchical model where MPI manages coarse-grained parallelism in non-shared memory systems. At a lower level, OpenMP processes within each MPI process manage fine-grained parallelism taking advantage of the shared memory to improve performance.

The main contributions of VFFVA are the complete use of C, which impacted mainly the computing time of small models (*n* < 3000) and the dynamic load balancing that was the main speedup factor for larger models.

### Impact on computing small models

VFFVA and FFVA were run five times on small models, i.e., Ecoli_core, EcoliK12, P_putida. VFFVA had at least 20 fold speedup (Table 2). The main contributing factor was the use of C over MATLAB in all steps of the analysis. In particular, the loading time of MATLAB java machine and the assignment of workers through `parpool` was much greater than the analysis time itself.

**Table 2:** Comparison of run times of FFVA and VFFVA in small models in seconds.

| Model | FFVA<br>mean(std)<br>loading and<br>analysis<br>time | VFFVA<br>mean(std)<br>loading and<br>analysis<br>time | FFVA<br>mean(std)<br>analysis<br>only |
| --- | --- | --- | --- |
| 2 cores |  |  |  |
| Ecoli_core | 19.5(0.5) | 0.2(0.01) | 0.37(0.1) |
| P_putida | 19.2(0.7) | 0.6(0.02) | 0.81(0.09) |
| EcoliK12 | 20.4(0.6) | 2.2(0.06) | 2.41(0.09) |
| 4 cores |  |  |  |
| Ecoli_core | 19.6(0.6) | 0.2(0.005) | 0.32(0.01) |
| P_putida | 19.4(1) | 0.5(0.02) | 0.61(0.01) |
| EcoliK12 | 20(0.8) | 1.3(0.04) | 1.64(0.08) |
| 8 cores |  |  |  |
| Ecoli_core | 19.4(0.5) | 0.2(0.03) | 0.35(0.05) |
| P_putida | 19.6(0.7) | 0.4(0.04) | 0.53(0.009) |
| EcoliK12 | 20(0.49) | 0.9(0.01) | 1.22(0.08) |
| 16 cores |  |  |  |
| Ecoli_core | 20.2(0.4) | 0.2(0.008) | 0.41(0.05) |
| P_putida | 19.5(0.4) | 0.4(0.04) | 0.51(0.03) |
| EcoliK12 | 22(0.7) | 0.7(0.01) | 0.87(0.03) |
| 32 cores |  |  |  |
| Ecoli_core | 22.2(0.4) | 0.3(0.008) | 0.6(0.12) |
| P_putida | 21.5(0.6) | 0.4(0.01) | 0.53(0.004) |
| EcoliK12 | 21.5(0.6) | 0.6(0.03) | 0.78(0.04) |

The result highlighted the power of C in gaining computing speed, through managing the different low-level aspects of memory allocation and variable declaration.

In the analysis of large models, where MATLAB loading time becomes less significant, dynamic load balancing becomes the main driving factor of the gained speedup.

### Impact on computing large models

The speedup gained on computing large models (Recon2 and E_Matrix) reached three folds with VFFVA (Figure 2) at 32 threads with Recon 2 (35.17*s* vs 10.3*s*) and E_Matrix (44*s* vs 14.7*s*). In fact, with dynamic load balancing, VFFVA allowed to update the assigned chunks of iterations to every worker dynamically, which guarantees an equal distribution of the load. In this case, the workers that get fast-solving LPs, will get a larger number of iterations assigned. Conversly, the workers that get ill-conditioned LPs and require more time to solve them, will get fewer LPs in total. Finally, all the workers synchronize at the same time to reduce the results. Particularly, the speedup achieved with VFFVA increased with the size of the models and the number of threads (Figure 2-E_Matrix). Finally, the different load balancing strategies (static, guided and dynamic) were compared further with two of the largest models (E_c__Matrix and Harvey).

**Figure 2:**
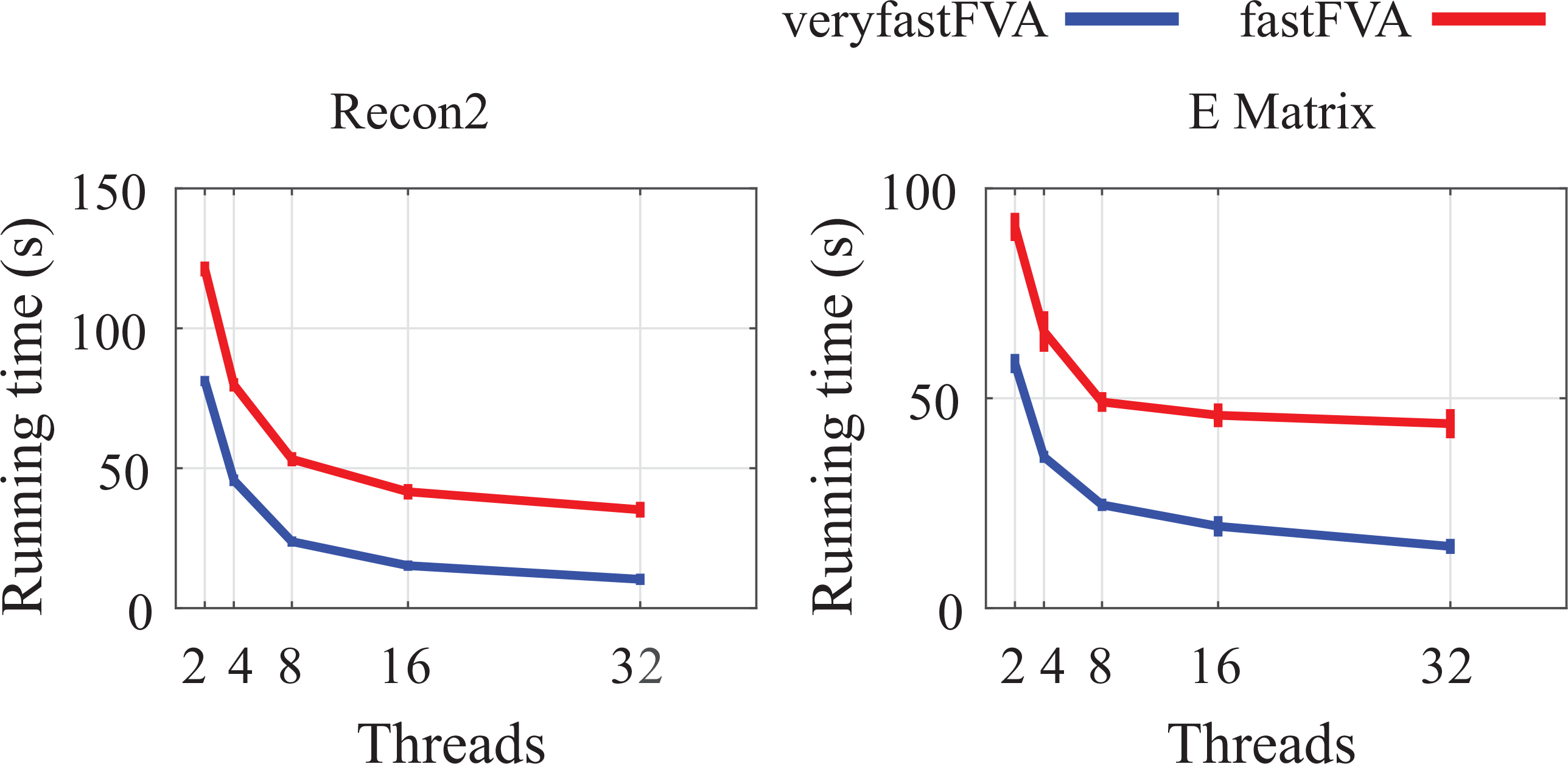
Run times of Recon2 and E_Matrix model using FFVA and VFFVA on 2,4,8,16, and 32 threads. The guided schedule was used in the benchmarking.

### Load management

Load management describes the different approaches to assign iterations to the workers. It can be static, where an even number of iterations is assigned to each worker. Guided schedule refers to dividing the iterations in chunks of size *n/workers* initially and *remaining_iterations/workers* afterward. The difference with static lies in the dynamic assignment of chunks, in a way that fast workers can process more iteration blocks. Finally, the dynamic schedule is very similar to guided except that chunk size is given by the user, which allows greater flexibility. In the following section, the load balancing strategies of E_c__Matrix and Harvey models were compared.

### Static schedule

Using static schedule, VFFVA assigned an equal number of iterations to every worker. With 16 threads, the number of iterations per worker equaled 1715 and 1716 (Figure 3-C). Expectedly, the run time varied widely between workers (Figure 3-B) and resulted in a final time of 393*s*.

**Figure 3:**
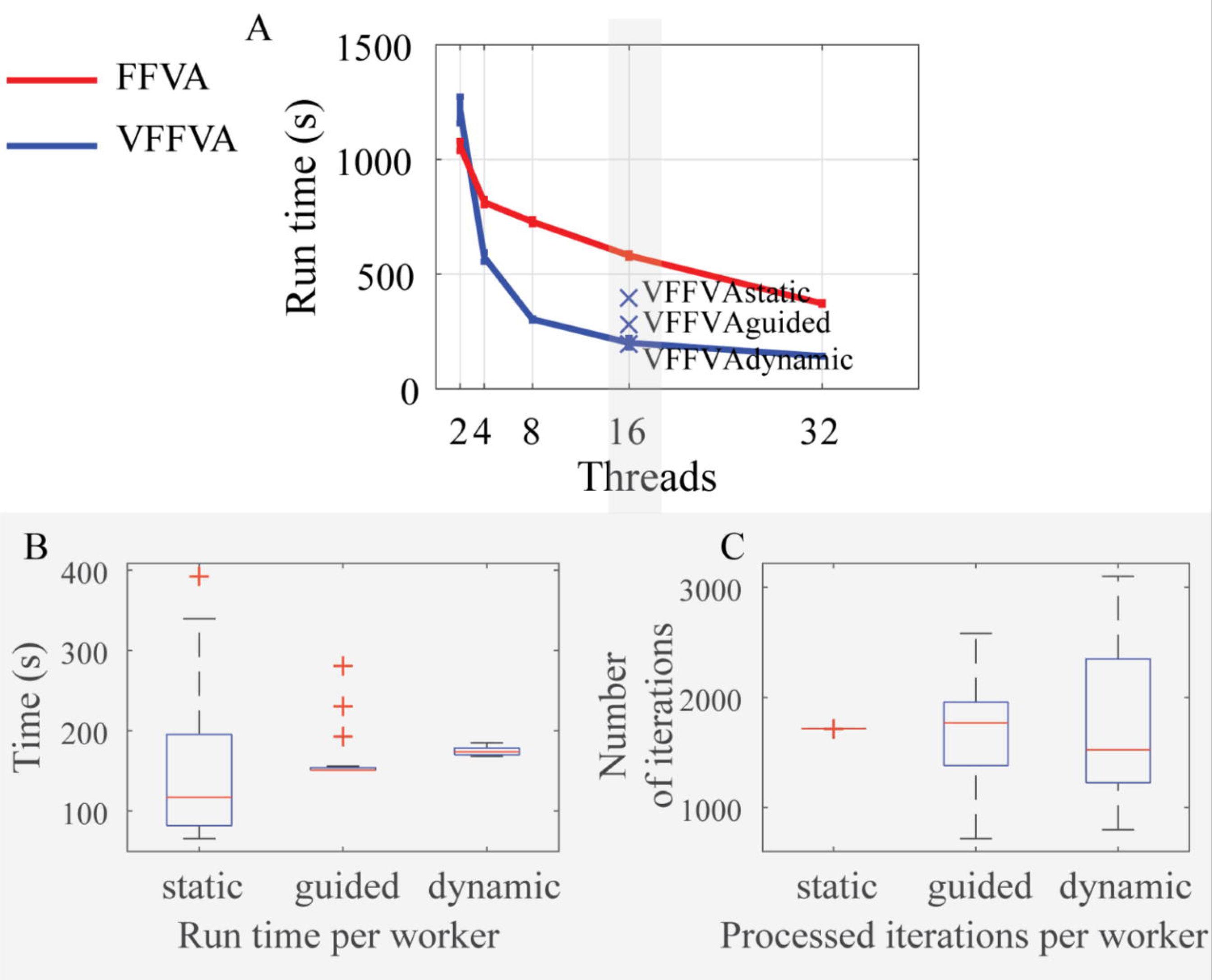
Run times of E_c__Matrix model. A-Run times of E_c__Matrix model at 2,4,8,16, and 32 threads using FFVA and VFFVA. B-Run time per worker in the static, guided, and dynamic schedule using 16 threads. C-The number of iterations processed per worker in the static, guided, and dynamic schedule using 16 threads.

### Guided schedule

With guided schedule (Figure 3-A), the highest speedup (2.9) was achieved with 16 threads (Figure 3-B). The run time per worker was quite comparable, and the iterations processed varied between 719 and 2581. The final run time was 281*s*.

### Dynamic schedule

Using a dynamic load balancing with a chunk size of 50 resulted in similar results to the guided schedule. The final run time equaled 197*s*, while FFVA took 581*s*. An optimal chunk size has to be small enough to ensure a frequent update on the workers’load, and big enough to take advantage of the solution basis reuse in every worker. At a chunk size of one, i.e., each worker is assigned one iteration at a time, the final solution time equaled 272*s*. In fact, for a small chunk size, the worker is updated often with new pieces of iterations, looses the stored solution basis of the previous problem, and has to solve the LP from scratch which slows the overall process.

Similarly, Harvey *Homo sapiens* metabolic model [24] (Figure 4-A) had a 2-fold speedup with 16 threads using a chunk size of 50 (806 mn) compared to FFVA (1611 mn). The run times with guided schedule (905 mn), dynamic schedule with chunk size 100 (850 mn) and chunk size 500 (851 mn) were less efficient due to the slower update rate leading to a variable analysis time per worker (Figure 4-B,C,D). VFFVA on eight threads (1323 mn with chunk size 50) proved comparable to FFVA (1214 mn) and distributedFBA (1182 mn) on 16 threads, thereby saving computational resources and time.

**Figure 4:**
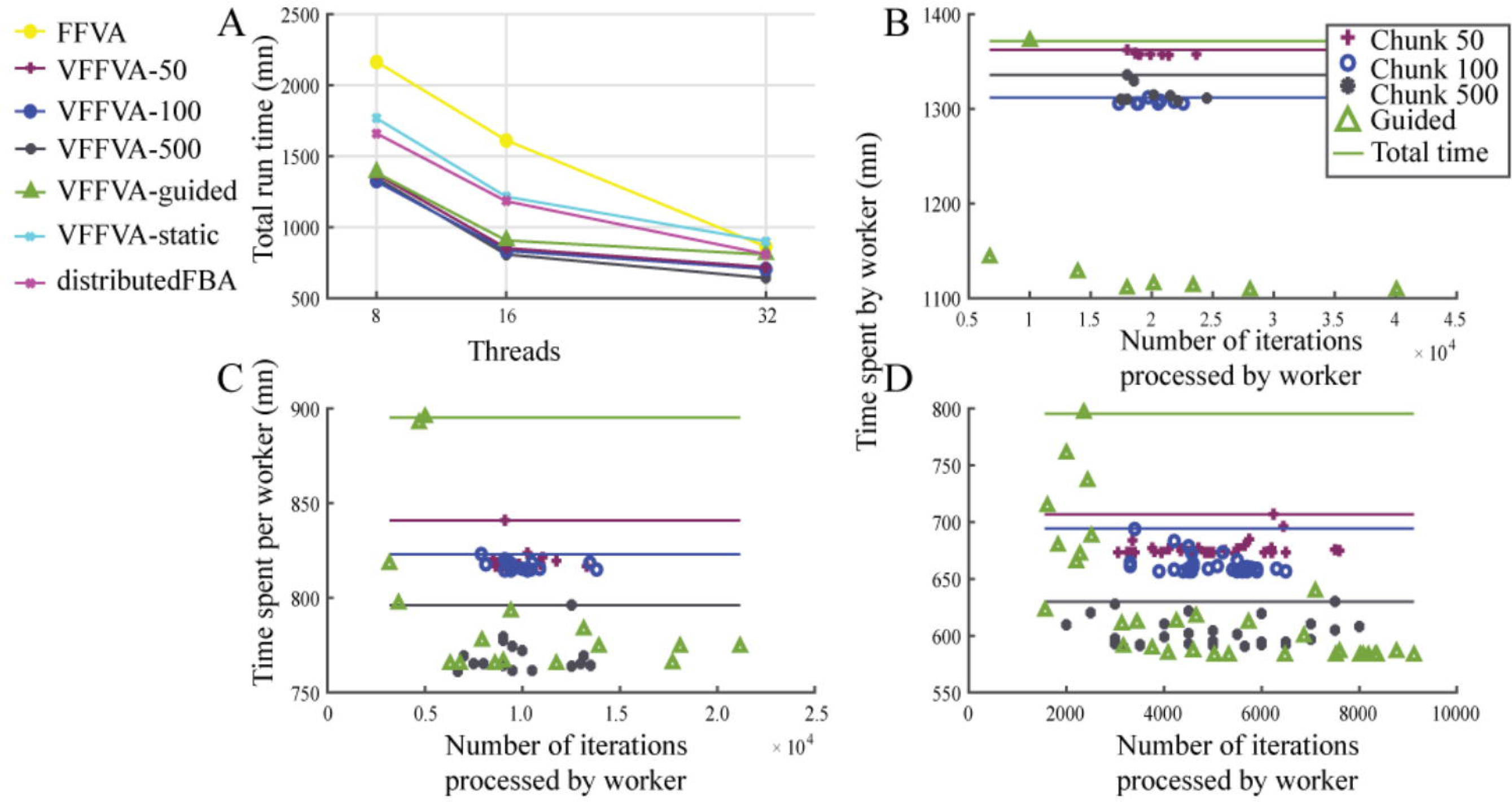
Run times per worker of Harvey *Homo sapiens* metabolic model. A-Total run time of the different load balancing schedules at 8, 16, and 32 threads. B-Run time per worker as a function of the number of iterations processed using the guided schedule and the dynamic schedule with a chunk size of 50, 100, and 500 with eight threads, C-16 threads, and D-32 threads.

### Impact on memory usage

In MATLAB, the execution of *j* parallel jobs implies launching *j* instances of MATLAB. On average, one instance needs 2 Gb. In a parallel setting, the memory requirements are at a minimum 2*j* Gb, which can limit the execution of highly parallel jobs. In the Julia-based distributedFBA, the overall memory requirement exceeded 15 Gb at 32 cores. VFFVA requires only the memory necessary to load *j* instances of the input model, which corresponds to the MPI processes as the OpenMP threads save additional memory through sharing one instance of the model. The differences between the FFVA and VFFVA get more pronounced as the number of threads increases (Figure 5) i.e., 13.5 fold at eight threads, 14.2 fold at 16 threads, and 14.7 fold at 32 threads.

**Figure 5:**
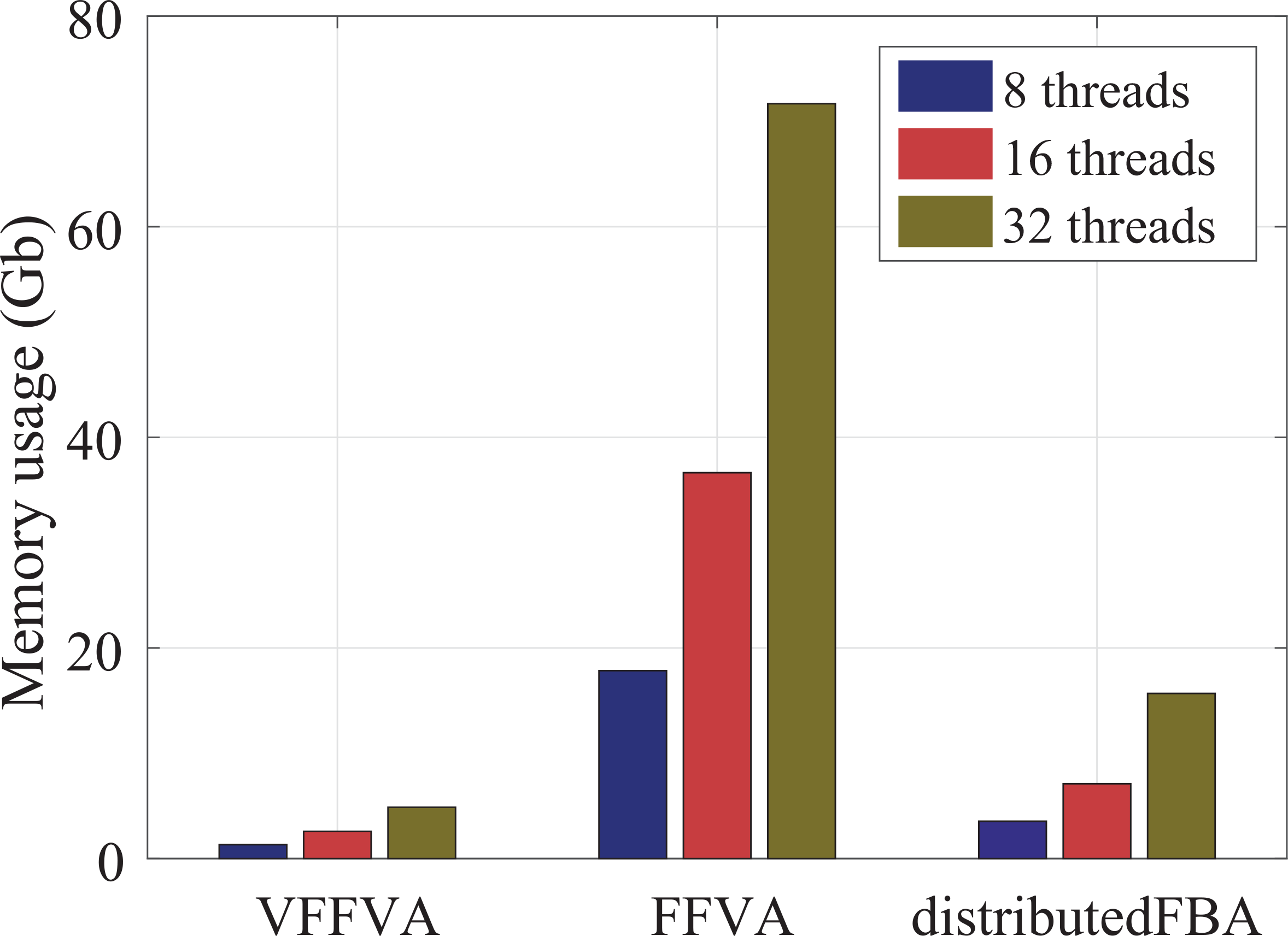
Physical memory usage at 8, 16, and 32 threads using FFVA, VFFVA, and dis-tributedFBA highlights a lower memory usage with VFFVA.

Finally, VFFVA outran FFVA and distributedFBA both on execution time and memory requirements (Table 3). The advantage becomes important with larger models and a higher number of threads, which makes VFFVA particularly suited for analyzing the exponentially-growing-in-size metabolic models in HPC setting.

**Table 3:** Comparative summary of the methods’ features.

| Feature | VFFVA | distributedFBA | FFVA | FVA |
| --- | --- | --- | --- | --- |
| Speed | ++++ | +++ | ++ | + |
| Memory | +++ | ++ | + | + |
| Load balancing | dynamic | static | static | static |

### Creation of warmup points for sampling

The generation of 30,000 warmup points were compared using the COBRA toolbox function createWarmup_MATLAB_ and a dynamically load-balanced C implementation createWarmup_VF_ on a set of models (Table 4). Since the COBRA toolbox implementation does not support parallelism, it was run on a single core and divided the run time by the number of cores to obtain an optimistic approximation of the parallel run times. The speedup achieved varied between four up to a factor of 100 in the different models (Table 4). Similarly to FFVA [7], the main factor for the speedup was the C implementation that allowed the reuse of the LP object in every iteration and save presolve time. Equally, the dynamic load balancing between workers ensured a fast convergence time.

Taken together, the dynamic load balancing strategy allows the efficient parallel solving of metabolic models through accelerating the computation of FVA and the fast preprocessing of sampling points thereby enabling the modeller to tackle large-scale metabolic models.

**Table 4:**
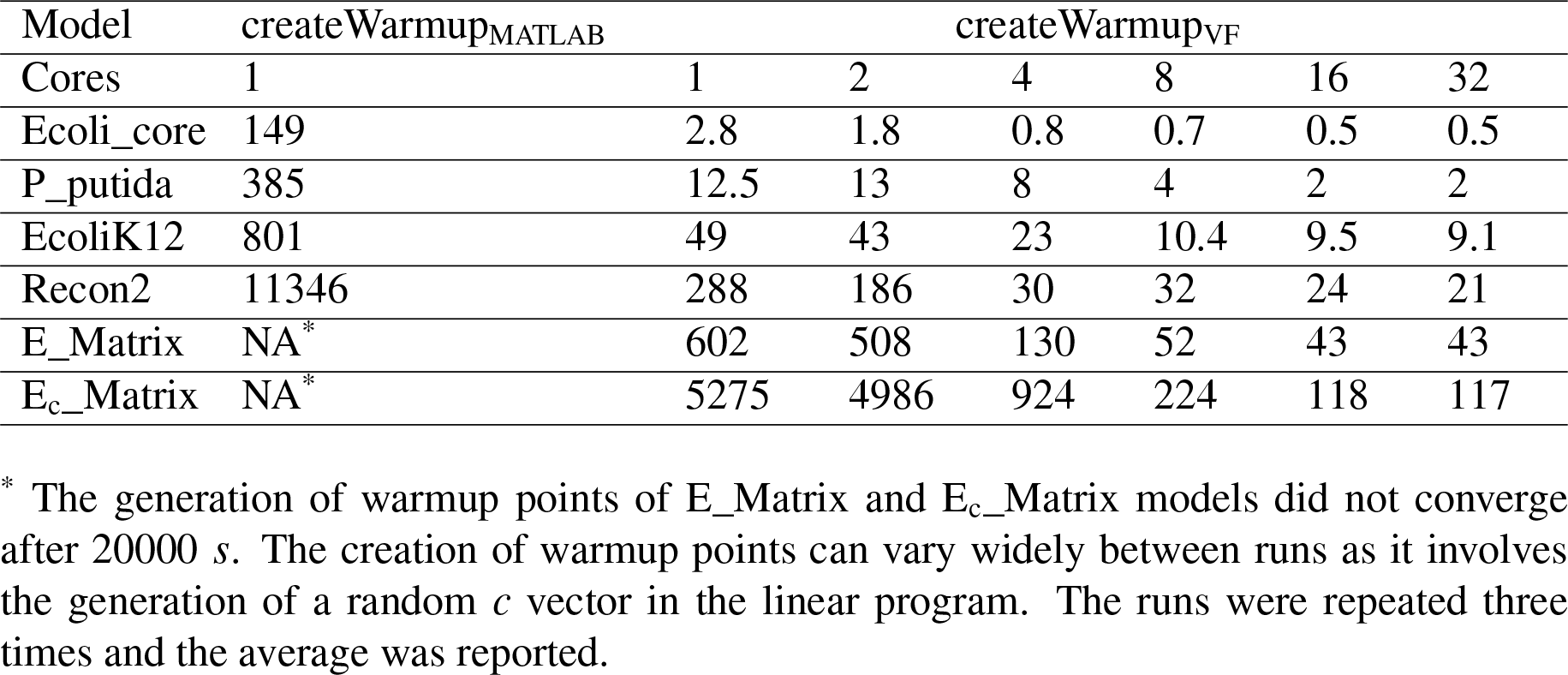
Generation of sampling warmup points using dynamic load balancing.

| Model | `createWarmup<sub>MATLAB</sub>` | `createWarmup<sub>VF</sub>` |  |  |  |  |  |
| --- | --- | --- | --- | --- | --- | --- | --- |
| Cores | 1 | 1 | 2 | 4 | 8 | 16 | 32 |
| Ecoli_core | 149 | 2.8 | 1.8 | 0.8 | 0.7 | 0.5 | 0.5 |
| P_putida | 385 | 12.5 | 13 | 8 | 4 | 2 | 2 |
| EcoliK12 | 801 | 49 | 43 | 23 | 10.4 | 9.5 | 9.1 |
| Recon2 | 11346 | 288 | 186 | 30 | 32 | 24 | 21 |
| E_Matrix | NA* | 602 | 508 | 130 | 52 | 43 | 43 |
| E<sub>c</sub>_Matrix | NA* | 5275 | 4986 | 924 | 224 | 118 | 117 |

\* The generation of warmup points of E\_Matrix and E<sub>c</sub>\_Matrix models did not converge after 20000 *s*. The creation of warmup points can vary widely between runs as it involves the generation of a random *c* vector in the linear program. The runs were repeated three times and the average was reported.

## Acknowledgements

The author would like to thank Valentin Plugaru at the University of Luxembourg for useful comments and guidance as well as fastFVA authors for publicly sharing their code and IBM for providing a free academic version of ILOG CPLEX. The experiments presented in this paper were partly carried out using the HPC facilities of the University of Luxembourg [25] - see http://hpc.uni.lu.

